# The day/night difference in the circadian clock’s response to acute lipopolysaccharide and the rhythmic Stat3 expression in the rat suprachiasmatic nucleus

**DOI:** 10.1101/342568

**Authors:** Simona Moravcová, Dominika Pačesová, Barbora Melkes, Hana Kyclerová, Veronika Spišská, Jiří Novotný, Zdeňka Bendová

## Abstract

The circadian clock in the suprachiasmatic nucleus (SCN) regulates daily rhythms in physiology and behaviour and is an important part of the mammalian homeostatic system. Previously, we have shown that systemic inflammatory stimulation with lipopolysaccharide (LPS) induced the daytime-dependent phosphorylation of STAT3 in the SCN. Here, we demonstrate the LPS-induced *Stat3* mRNA expression in the SCN and show also the circadian rhythm in *Stat3* expression in the SCN, with high levels during the day. Moreover, we examined the effects of LPS (1mg/kg), applied either during the day or the night, on the rhythm in locomotor activity of male Wistar rats. We observed that recovery of normal locomotor activity patterns took longer when the animals were injected during the night. The clock genes *Per1, Per2* and *Nr1d1*, and phosphorylation of kinases ERK1/2 and GSK3β are sensitive to external cues and function as the molecular entry for external signals into the circadian clockwork. We also studied the immediate changes in these clock genes expressions and the phosphorylation of ERK1/2 and GSK3β in the suprachiasmatic nucleus in response to daytime or night-time inflammatory stimulation. We revealed mild and transient changes with respect to the controls. Our data stress the role of STAT3 in the circadian clock response to the LPS and provide further evidence of the interaction between the circadian clock and immune system.

## Introduction

The temporal organisation of behavioural and physiological functions in mammals mostly depends on the timing signals generated by the circadian clock in the suprachiasmatic nucleus (SCN). The SCN integrates external timing cues with an endogenous molecular clockwork and synchronises circadian oscillations in other brain parts and peripheral tissues (Albrecht, 2012). The morphology of the SCN varies across species, but its basic structure is shared by all mammals. It consists of two main parts: the shell or the dorsomedial part (dmSCN), which contains intrinsically rhythmic cells, and the core or the ventrolateral part (vlSCN), which receives most of the regulatory inputs from other parts of the brain (Morin, 2007). The molecular basis of a circadian clock involves several interlocking transcriptional loops of clock genes, such as *Clock*, *Bmal1*, *Period 1* (*Per1*)*, Per2, Cryptochrome 1* (*Cry1*) and *Cry2, Casein kinase 1 epsilon* (*CK1ε*)*, RevErbα* (*Nr1d1*) and *Rorα*. The stability of these loops is supported by posttranslational modifications of clock protein by several kinases, including glycogen synthase kinase-3beta (GSK3β) and p42/44 mitogen-activated protein kinase (ERK1/2) (Iitaka et al., 2005; Kurabayashi et al., 2006; Sanada et al., 2002; 2004; Reischl and Kramer, 2011). This kinase also plays a role in photic entrainment of the circadian clock in the SCN (Obrietan et al., 1998; Dziema et al., 2003); thus, it is an integral part of the circadian clock in the SCN.

Many studies have demonstrated that the susceptibility of organisms to inflammatory stimuli is strongly influenced by the SCN and that the immune responses differ in their strength, depending not only on the dose of the pathogen but also on the time of infection. Recently, Kiessling et al. (2017) reported reduced footpad swelling and the lowest immune cell recruitment when mice were infected with Leishmania parasitic during the early subjective day, compared to the late day or even the subjective night. This difference was apparently driven by a daytime-dependent neutrophil and anti-inflammatory macrophage infiltration to the infection site, which was regulated by the circadian clock. Furthermore, lipopolysaccharide (LPS) administration induces higher levels of proinflammatory cytokines and chemokines in a serum if mice are injected at the transition point between their rest and activity phases of the day than between the activity and rest phases at the beginning of the light phase of the day (Gibbs et al., 2012; Guerrero-Vargas et al., 2014). The circadian regulation of immune functions is also supported by the observations that the dysfunction of the timing system by either ablation of the SCN or by forced desynchronization severely perturbs the immune system. The SCN lesion enhances the level of LPS-induced corticosterone (Kalsbeek et al., 2012), modulates the temperature response to the LPS and markedly enhances tumour necrosis factor-α (TNF-α) and interleukin (IL-6) plasma levels after LPS treatment (Wachulec et al., 1997; Guerrero-Vargas et al., 2014). Mice exposed to chronic jet lag showed increased LPS-induced mortality compared to non-shifted controls (Castanon-Cervantes et al., 2010).

The current research also focuses on the reciprocal relationship between the immune system and the circadian clock. It has been shown that the LPS upregulates the p65 subunit of NF-κB transcription complex and immediate early-gene *c-Fos* in the SCN (Marpegán et al., 2005; Sadki et al., 2007; Beynon and Coogan, 2010). Other reports have suggested that the LPS-induced circadian responses are mediated by Toll-like receptor 4 (TLR4) (Paladino et al., 2010) and that the TNF-α and chemokine CCL2 mediate the signalling of peripheral LPS application into the SCN via changes of molecular clock in SCN astrocytes (Paladino et al., 2014; Duhart et al., 2013; 2016). The inflammatory signalling pathways in the brain mostly converge on the activation of signal transducers and activators of transcription 3 (STAT3) (Rummel, 2016). In our previous study, we demonstrated the daytime-dependent LPS-induced phosphorylation of STAT3 on Tyr705 and Ser727 in the SCN astrocytes (Moravcová et al., 2016). In the present study, we examine the daytime-dependent effect of peripheral LPS on the expression of the *Stat3* gene in the SCN. To detect a possible impact on the clockwork mechanisms, we assessed the expression of clock genes *Per1, Per2* and *Nr1d1*, which are sensitive to external cues and function as the molecular entry for external signals into the circadian clockwork. The phosphorylated forms of ERK1/2 (pERK1/2) and GSK3β (pGSK3β) play roles in stabilizing the clockwork mechanism, show specific expression patterns in the SCN and time-of-day-dependent response to morphine (Pačesová et al., 2015) and are even involved in STAT3 signalling (Beurel and Jope, 2008, 2009). Therefore, we also detected their response to LPS administration in the SCN by immunohistochemistry. We initially measured locomotor activity in our study to control for overall sickness after the LPS treatment. Interestingly, the rate of the physical recovery over the following days differed according to the time of LPS administration.

## Materials and methods

### Animals

Male Wistar rats (Velaz Ltd., Koleč, Czech Republic) were maintained under a 12-hr light-dark regimen (with lights on from 06:00 to 18:00) at a temperature of 23 °C ± 2 °C with free access to food and water for at least 2 weeks before the experiment. This study was carried out in strict accordance with the recommendations in the Guide for the Care and Use of Laboratory Animals of the National Institutes of Health. The protocol was approved by the Animal Protection Law of the Czech Republic (Protocol Number: MSMT-23852/2014-14). All sacrification was performed by rapid decapitation under sodium pentobarbital anesthesia, and all efforts were made to minimize suffering.

### Locomotor activity measurement

The locomotor activity of 20 rats was monitored using infrared motion detectors (Mini-Mitter VitalView data acquisition system) for three days before the experiment, which served as an internal control for total activity, acrophase and amplitude. On the day of the experiment, 10 rats received an intraperitoneal injection of LPS (from *E. coli*, strain 055:B5; 1 mg/kg; Sigma Aldrich) at ZT6, as did 10 rats at ZT15. Activity counts accumulated over a 1-h period were fitted with single cosine curves, as described by Soták et al. (2011) and Hahnová et al. (2016) and were analysed with GraphPad Prism version 6.00. The acrophase and amplitude were indicated from cosine waves. The data were plotted as the mean of 10 animals.

### Experimental design

#### Circadian rhythm of *Stat3* mRNA

Adult male rats were released into constant darkness at the time of dark-to-light transition (designated as circadian time 0; CT0). During the first cycle in darkness, the animals were sacrificed by rapid decapitation at 3-h intervals (n = 3-4); their brains were frozen on dry ice and stored at -80 °C.

#### Effect of LPS on levels of *Stat3* and clock genes mRNA, pERK1/2 and pGSK3β

Adult rats received an intraperitoneal injection of LPS (1 mg/kg) at ZT 6 or ZT 15. Time was expressed as Zeitgeber time (ZT), with ZT0 corresponding to the time of lights on and ZT12 corresponding to the time of lights off. The control animals received saline. Four experimental and four control animals were anesthetized with thiopental 2, 5, 8 and 24 hr later and were either killed by rapid decapitation for gene expression assessment by in situ hybridization or perfused through the ascending aorta with 4% paraformaldehyde in PBS, as described before (Bendová et al., 2012), for immunohistochemical detection of pERK1/2 and pGSK3β.

### *In situ* hybridisation

The cDNA fragments of rat *Stat3, Per1, Per2* and *Nr1d1* were used as templates for the in vitro transcription of complementary RNA probes (SP6/T7 MAXIscript kit, Applied Biosystems, Austin, TX, USA). The probes were labelled by [α-35S]-UTP (American Radiolabeled Chemicals, Inc., St. Louis, MO, USA) and purified using Chroma-Spin 100-DEPC H2O columns (Clontech Laboratories Inc., Mountain View, USA). In situ hybridisation was performed as described by Matějů et al. (2009). For each gene, brain sections from control and experimental rats were processed simultaneously under identical conditions. Autoradiographs were analysed using NIH Image J software to detect the relative optical density (OD) of the specific hybridisation signal. In each animal, the signal was quantified bilaterally at the mid-caudal SCN section. Each measurement was corrected for nonspecific background by subtracting the OD values from the adjacent area in the hypothalamus with a consistently low OD. The OD values for each animal were calculated as a mean of values for the left and right SCN.

### Quantitative Real Time RT-PCR

The brains were sectioned into a series of 20-μm-thick coronal slices throughout the rostral-caudal extent of the SCN. The slices were stained with ethanolic cresyl violet for 60 sec, and SCN regions were isolated using laser microdissection (LMD 6000; Leica) and immediately homogenised in RLT buffer (RNeasy Plus Micro kit; Qiagen). Total RNA was extracted with the Rneasy Plus Micro Kit (Qiagen) according to the manufacturer’s instructions. We converted 1 μg of total RNA to cDNA using two-step Enhanced Avian Reverse Transcriptase eAMV RT (Sigma-Aldrich) according to the manufacturer’s instructions. TaqMan^®^ PreAmp Master Mix (Life Technologies) was used to pre-amplify small amounts of cDNA after reverse transcription. Samples of pre-amplified cDNA (1 μl) were amplified in 20 μl of PCR reaction mixture containing 5x HOT FIREPol^®^ Probe qPCR Mix Plus (Baria) plus TaqMan probes (Life Technologies) for rat *Stat3* gene (Rn00680715_m1) and housekeeping gene *Actb* (Rn00667869_m1). All of the qPCRs were performed in duplicate on a LightCycler^®^ 480 Instrument (Roche Life Science, Indianapolis, IN, USA) using the following temperature profile: initial denaturation at 95 °C for 15 min, followed by 60 cycles consisting of denaturation at 95 °C for 18 secs and annealing/elongation at 60°C for 60 sec. The mean of the crossing point (Cp) obtained from qPCR was normalised to the level of housekeeping gene *Actb* and then used to analyse relative gene expression by the ΔΔCT method (Livak and Schmittgen, 2001).

### Immunohistochemistry

The brains were sectioned into a series of 30-μm-thick free-floating coronal slices throughout the rostral-caudal extent of the SCN. The levels of phospho-p44/42 MAPK (Thr202/Tyr204) and pGSK3β (Ser9) (antibodies purchased from Cell Signalling Technology, Inc., Danvers, MA) were assessed by immunohistochemistry using the avidin/biotin method with diaminobenzidine as the chromogen (Vectastain ABC Kit, Vector, Burlingame, CA, USA). All brain sections were processed simultaneously under identical conditions. Immunopositive cells in the SCN mid-caudal region were manually tagged and counted using an image-analysis system (NIH Image J software). To delineate the position of the ventrolateral and dorsomedial SCN, the boundaries of the pERK1/2 signal at ZT15 and ZT6, respectively, were saved as regions of interest and applied to all immunohistochemical images. The data were expressed as the means of values from the left and right SCN.

### Statistical analysis

The data were expressed as the mean of the values from the left and right SCN and as the mean ± SEM of the number of animals per time point. The data were analysed by multiple t-tests, with the Sidak-Bonferroni method used for statistical significance. Two-way ANOVAs were performed to compare trends in the total activity, acrophase and amplitude of rhythms in the locomotor activity between groups. The circadian profile of *Stat3* mRNA levels and locomotor activity rhythmicity were analysed by a one-way ANOVA for the time differences. P < 0.05 was required for significance. Moreover, the circadian profiles were fitted with single cosine curves (Soták et al., 2011; Hahnová et al., 2016), defined by the equation: [Y= mesor+(amplitude*-cos(2*p*(X-acrophase)/wavelength] with a constant wavelength of 24 h.

## Results

### The recovery of locomotor activity from systemic LPS depends on the time of its administration

To monitor the recovery from acute infection in natural conditions, we measured the locomotor activity of rats for eight consecutive LD cycles following the systemic LPS administration (Fig. 1). The monitoring for three cycles before LPS treatment served as an internal control for each group (plotted only the last day before the treatment; Fig. 1A). During these days, the mean sum of activity counts per day (total activity) was 3632 for the rat group that was later treated with LPS at ZT6 and 3536 for the group that was later treated at ZT15 (Fig. 1B). The amplitude was 149.3 counts/h for the ZT6 group and 150 counts/h for the ZT15 group. The two-way ANOVA revealed a significant difference between the activity cycles of both groups in the first day after their LPS treatment (LD1; F (1, 445) = 22.31; P < 0.0001), second day (LD2; F (1, 450) = 6.302; P = 0.0124), third day (LD3; F (1, 450) = 19.14; P < 0.0001) and fourth day (LD4; F (1, 450) = 16.7; P < 0.0001) (Fig. 1A). One-way ANOVA confirmed the significant rhythmicity in both groups already on LD1 (ZT6 group, F (24, 213) = 21.31; P < 0.0001; ZT15 group, F (24, 213) = 3.624; P < 0.0001). Although the two-way ANOVA did not confirm the difference between groups in total activity, the pairwise comparisons revealed a significant difference between total activity values on LD3 (P = 0.0014) (Fig. 1B). The two-way ANOVA revealed a significant difference between both groups in the gradual recovery of amplitude (F (1,8) = 10.56; P = 0.0117) and thus confirmed that the recovery of locomotor activity was faster in animals treated with LPS during the day than during the night.

**Fig. 1.**
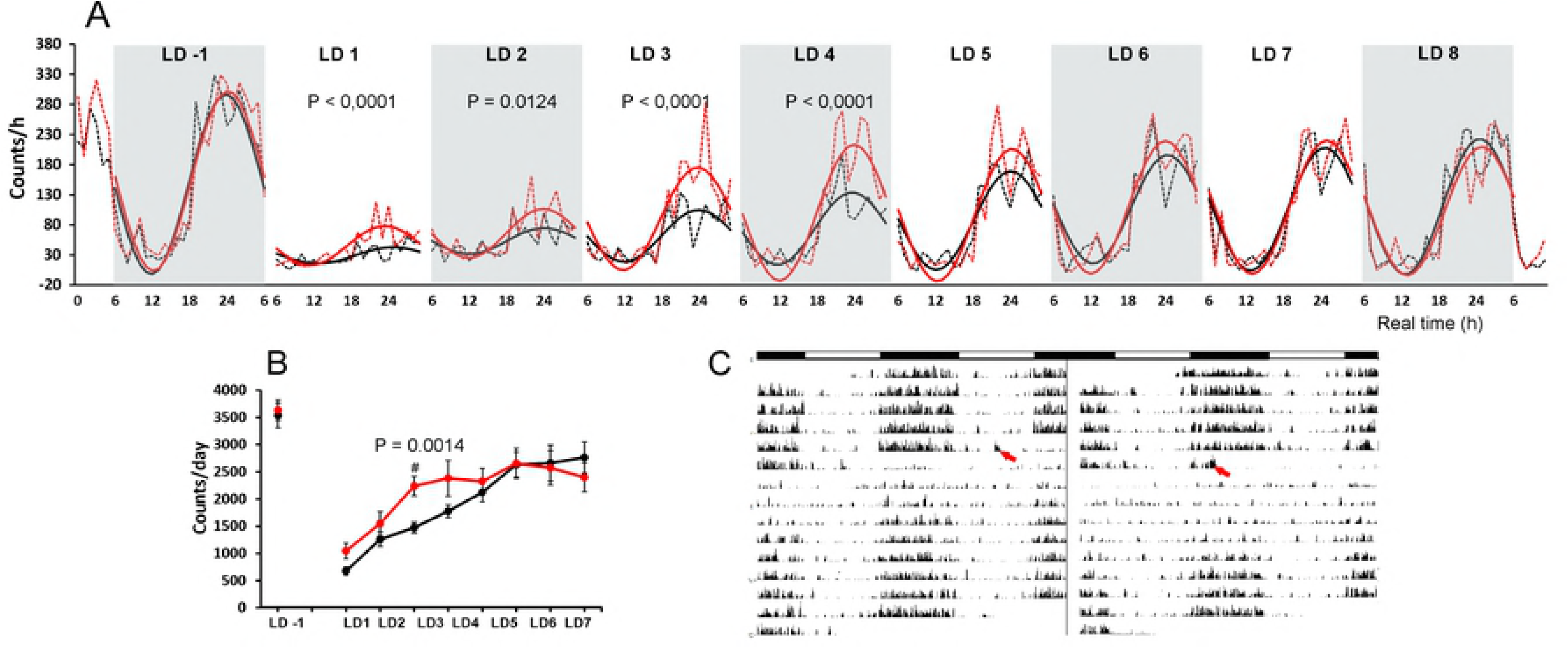
The recovery of normal locomotor activity patterns took longer when the animals were injected during the night. (A) The changes in locomotor activity one day before (LD-1) and 8 days after (LD1–LD8) the LPS (1 mg/kg) was applied at ZT6 (i.e., at real time (RT12); red line) or ZT15 (i.e., at real time 21 (RT21); black line) and administered at LD 0 (not plotted). The mean total activity counts during the 1-hr period were plotted (dotted lines) along with the population cosine wave for better resolution of the difference (full lines). The two-way ANOVA results for significance are shown. (B) The total activity counts per day, which show slower recovery by the animals injected at ZT15 compared to the animals injected at ZT6. (C) The representative actograms of animal injected at ZT6 (on the left) and at ZT15 (on the right). The times of LPS administration are indicated with red arrows.

### Circadian rhythmicity of *Stat3* expression and the time-dependent effect of acute LPS on its mRNA level in the rat SCN

We performed *in situ* hybridisation of the SCN sections to determine whether the mRNA expression of *Stat3* would show similar circadian rhythmicity as the STAT3 protein (Moravcová et al., 2016). One-way ANOVA revealed that under DD conditions, the *Stat3* mRNA expressed circadian rhythm in the SCN (P = 0.0012; Fig. 2). The cosinor analyses also confirmed a significant circadian rhythmicity (R2 = 0.5327, P < 0.0002).

**Fig. 2.**
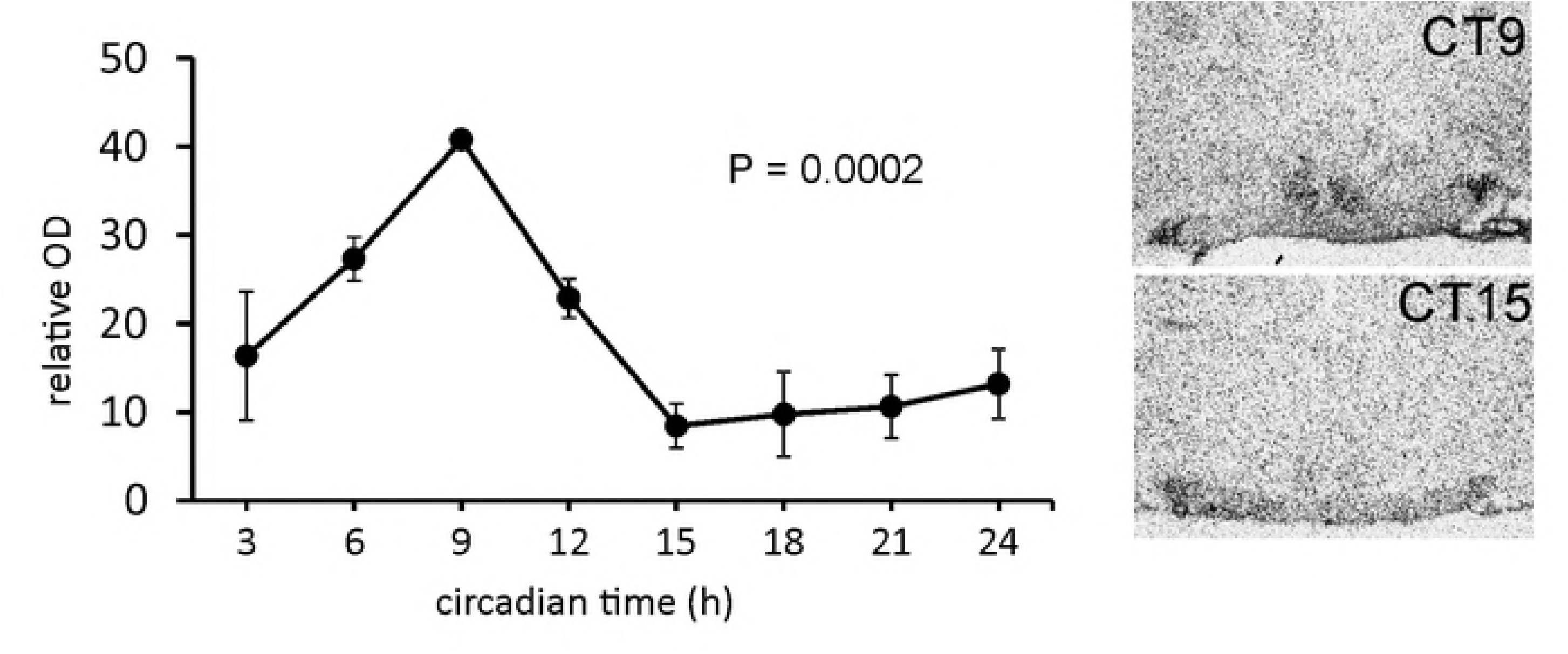
Rhythmic expression of *Stat3* mRNA in the rat SCN. The data are expressed as relative optical density values of the specific hybridisation signal. Each point represents the mean ± SEM from 3-4 animals. The P-value represents the significance of circadian rhythmicity revealed by the cosinor analyses. Representative photomicrographs of coronal sections of the SCN, probed for *Stat3* mRNA, which demonstrate the highest- and lowest-intensity hybridisation signal.

To reveal the effect of acute LPS on the *Stat3* expression, animals were injected with LPS at ZT6—the time of increasing *Stat3* endogenous expression—and at ZT15—the time of minimum *Stat3* expression in the SCN. We performed *in situ* hybridisation and revealed that when LPS was applied during the day, the level of *Stat3* mRNA did not change significantly until the 8 h from the injection (P = 0.0006), while when applied at ZT15, it markedly induced the *Stat3* transcription 2 h (P = 0.0004) and 5 h (P = 0.0056) after application (Fig. 3A). In the nightime, the LPS induced the *Stat3* mRNA of 11 ± 0.86 of the control values in 2 h, and 2.7 ± 0.37 times in 5 h (Fig. 3D). To verify the results obtained from *in situ* hybridisation, we checked the day/night difference in LPS effect at one time point using RT-PCR from the SCN isolated by laser dissection (Fig. 3B, C). Multiple t-tests with the Sidak-Bonferroni method showed that the mRNA level increased significantly after LPS at both ZT6 (P = 0.0091) and ZT15 (P = 0.0096). However, in the daytime, the LPS induced the *Stat3* mRNA of only 1.417 ± 0.1598 of the control values, but LPS applied at ZT15 increased the *Stat3* mRNA 15.8 ± 2.6 times (Fig. 3E). This result thus corresponds with the data obtain by *in situ* hybridization and confirms that the *Stat3* expression would show greater LPS-induced increase during the night apparently due to the low endogenous mRNA level during the night (Fig. 2). The significant difference between the day and night control groups (P < 0.0001; Fig. 3B) confirms the day/night difference of *Stat3* expression revealed by optic density measurements of *in situ* autoradiographs (Fig. 2).

**Fig. 3.**
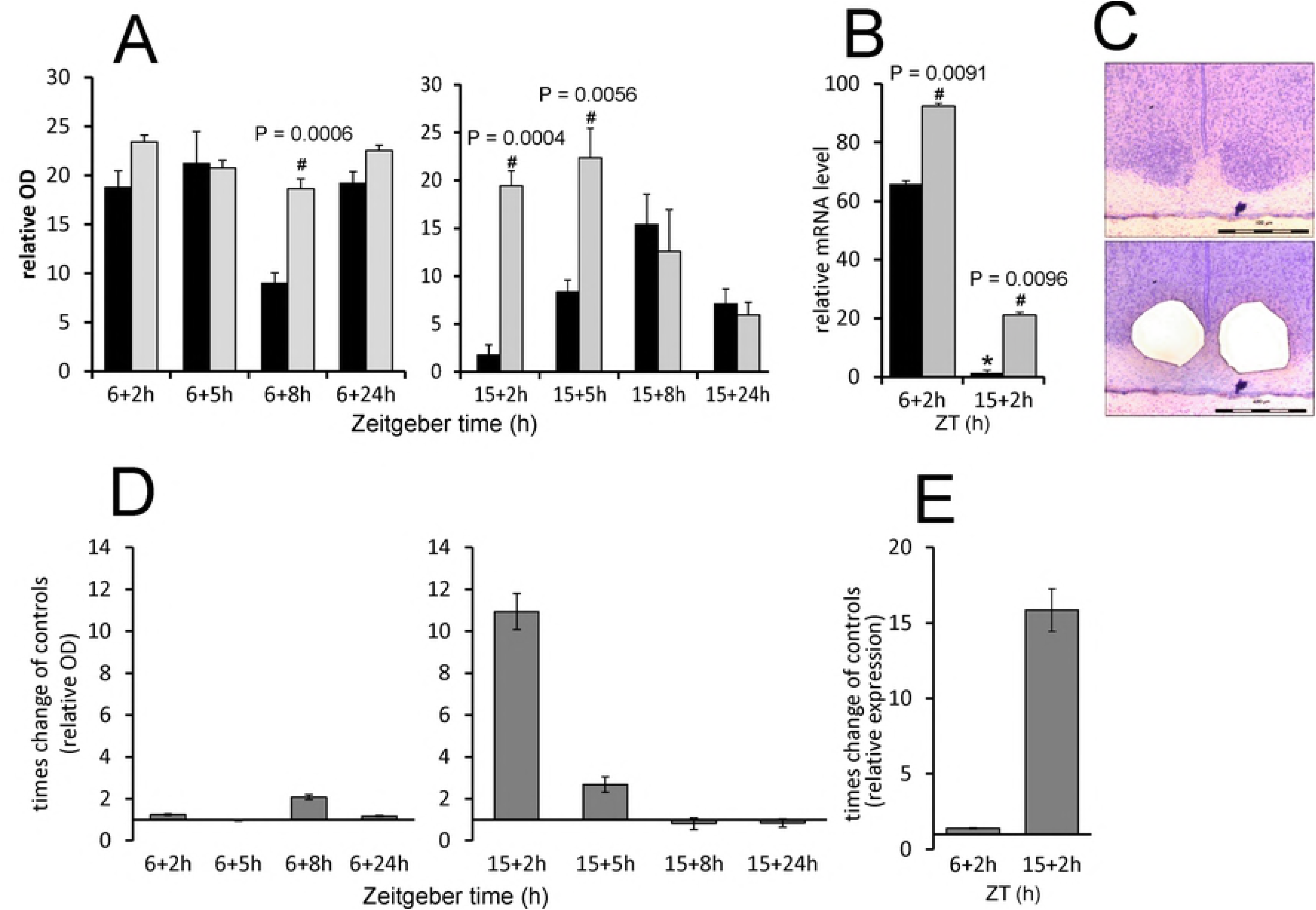
LPS-induced changes in *Stat3* gene expression in rat SCN. Adult rats were injected with LPS (1 mg/kg) either during the day at ZT6 or at night at ZT15 and were sampled 2 h, 5 h, 8 h and 24 h later (grey columns) (controls; black columns). (A) The levels of *Stat3* mRNA were assessed via the intensity of the hybridisation signal. Each column represents the mean of four values ± SEM. (B) The transcription level of *Stat3* was determined using quantitative RT-PCR, and the normalised values were converted to a percentage of the maximum value of the transcript. Each point represents the mean ± SEM of 3 animals. # P: The value of multiple t-tests with the Sidak-Bonferroni post-hoc test. (C) Representative photomicrographs of coronal sections of the SCN demonstrate the area of laser dissection of the SCN. Scale bar = 400 μm. (D, E) The data from A (B resp.) plotted as a LPS-induced times change in the *Stat3* expression compared to controls show the direct effect of LPS regardless of the daily gene oscillations.

### Time-dependent effect of acute LPS on the clock gene level in rat SCN

In the clockwork machinery, the genes *Per1, Per2* and *Nr1d1* best reflect the immediate changes in the external environment of the clock. To reveal whether the acute LPS may affect the clockwork mechanism distinctly depending on the time of day, we followed the changes in the expression of these clock genes in the SCN by *in situ* hybridisation. The clock genes show high-amplitude circadian rhythmicity in their expression. To extract the direct effect of LPS from the clock-controlled genes transcription, we plotted the data as the times difference between values from LPS-treated groups and controls. The real values are summarized in supplemental Fig.1. As shown in Fig. 4A, a significant reduction of *Per1* mRNA was observed 24 h after the daytime LPS application (P = 0.0024). A significant induction of *Per2* was observed 8 h after daytime LPS application (P = 0.0131) (Fig. 4B). No differences between the control groups and LPS-treated animals were detected for *Per1* and *Per2* expression in the SCN after the night-time application. In contrast, the level of *Nr1d1* was upregulated 8 h after the night-time LPS injection (P = 0.0019) but was not affected by the LPS applied at ZT6 (Fig. 4C).

**Fig. 4.**
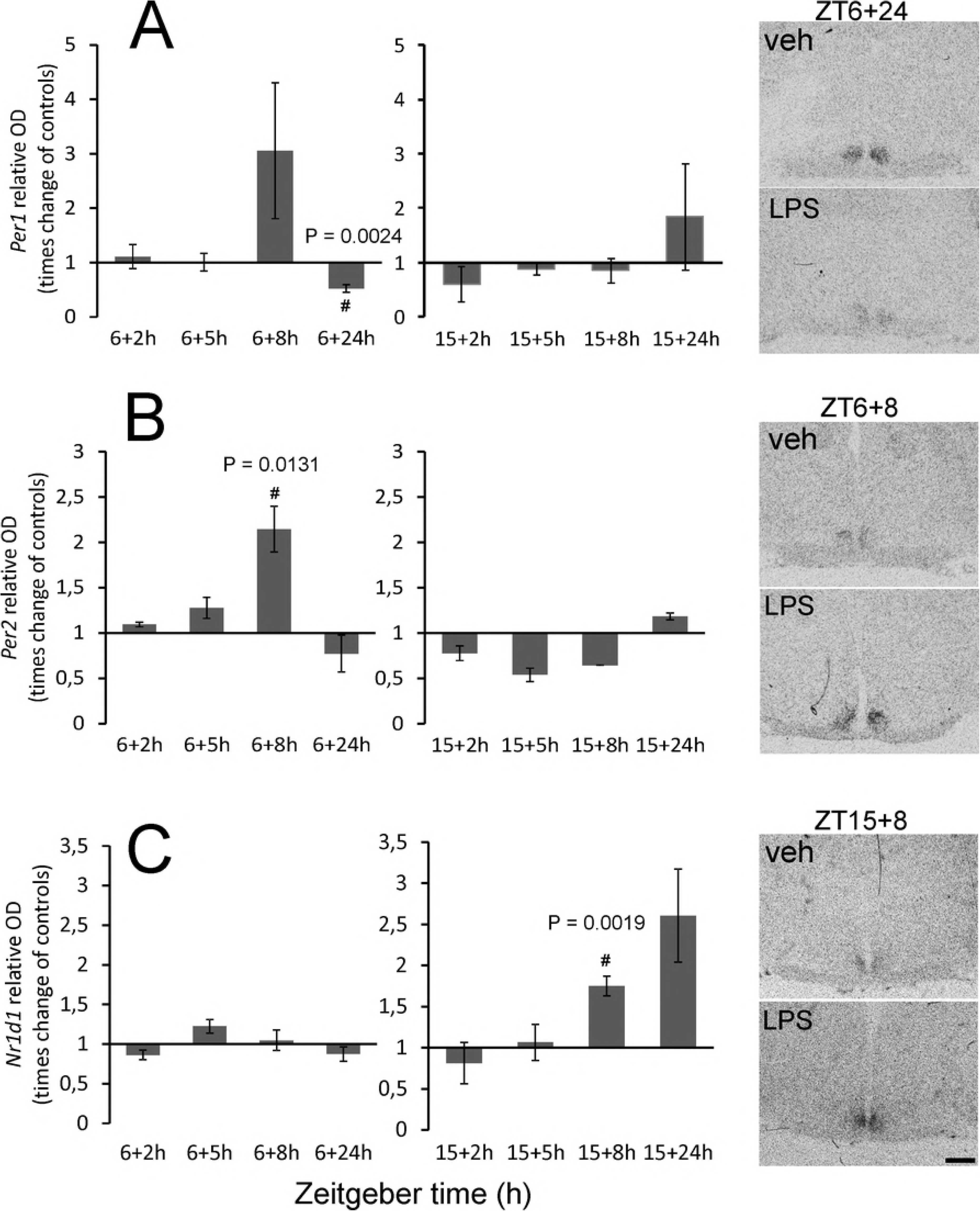
Effect of acute systemic LPS administration on clock gene expression in rat SCN. Adult rats were injected with LPS (1 mg/kg) either during the day, at ZT6, or at night, at ZT15, and sampled 2 h, 5 h, 8 h and 24 h later. The levels of *Per1* (A), *Per2* (B) and *Nr1d1* (C) mRNAs were assessed as the intensity of the hybridisation signal. Each column represents the mean of four values ± SEM. # P: Values of multiple t-tests with the Sidak-Bonferroni post-hoc test. The representative photomicrographs of coronal sections of the SCN demonstrate the intensity of the signals for each control/LPS pair that showed statistically significant differences. Scale bar = 500 μm.

### Time-dependent effect of acute LPS on pERK1/2 and pGSK3β levels in rat SCN

The phosphorylation state of both kinases shows significant circadian rhythmicity in the SCN. To better distinguish the effect of LPS from the clock-controlled baseline levels, we plotted the data as the times difference between values from LPS-treated groups and controls. The real values are summarized in supplemental Fig.2. The acute application of LPS at a dose of 1 mg/1kg showed opposite effects on pERK1/2 levels when applied during the day versus the night. LPS applied at ZT6 significantly induced pERK1/2 in the vlSCN within 2 h (P = 0.0190) (Fig. 5A) and in the dmSCN within 2 h (P = 0.0175) (Fig. 5B). LPS applied at ZT15 significantly reduced the number of pERK1/2 immunopositive cells in the vlSCN within 8 h (P = 0.0209) (Fig. 5A). There was no significant change in pERK1/2 levels in the dmSCN at night.

**Fig. 5.**
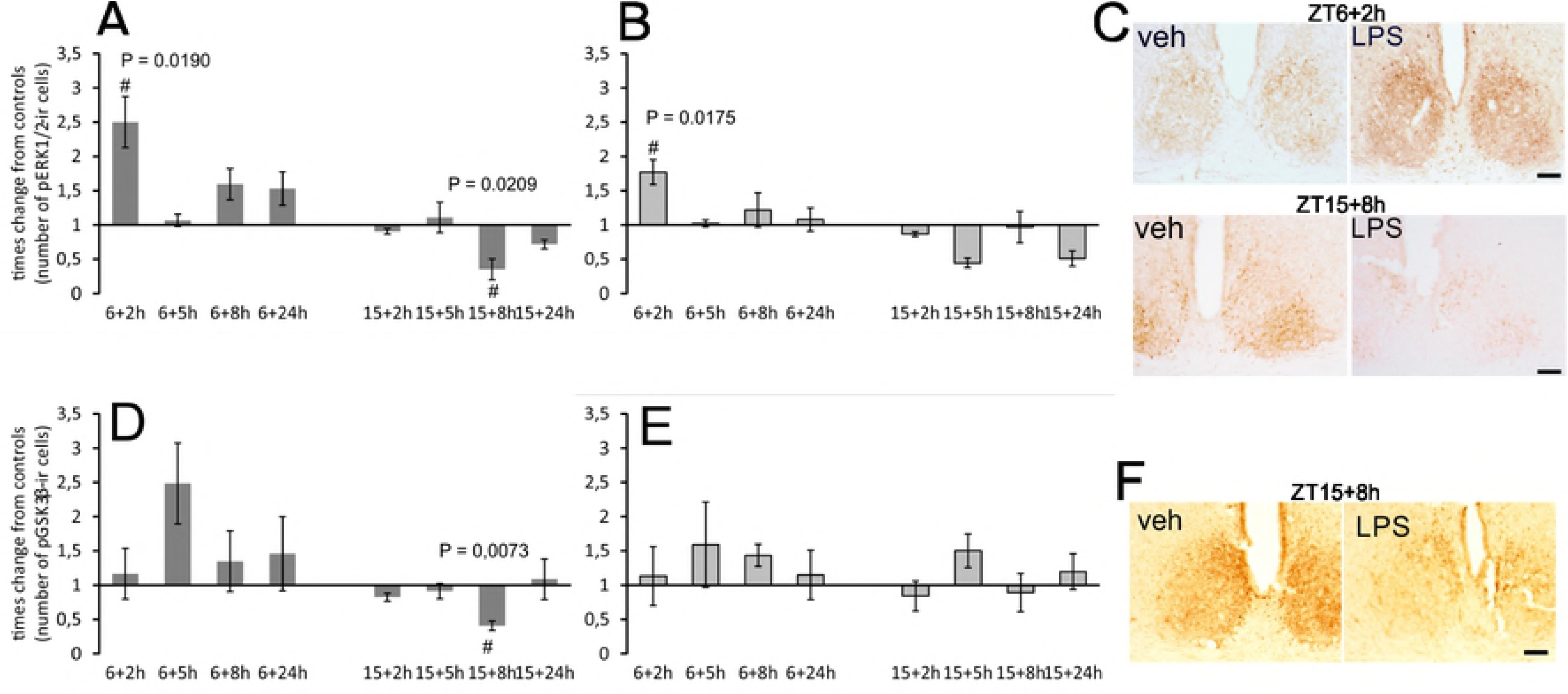
Effect of acute systemic LPS administration on ERK1/2 (A, B) and GSK3β (D, E) phosphorylation within rat SCN. Adult rats were injected with LPS (1 mg/kg) either during the day at ZT6 or at night at ZT15 and sampled 2 h, 5 h, 8 h and 24 h later. Levels of immunopositive cells were assessed separately for the ventrolateral (A, D) and dorsomedial (B, E) SCN. Each column represents the mean of four values ± SEM. # P: Values of multiple t-tests with the Sidak-Bonferroni post-hoc test. The representative photomicrographs of coronal sections of the SCN demonstrate the intensity and distribution of pERK1/2 (C) and p GSK3β (F) in the control and LPS-treated animals when the control/LPS pairs showed statistically significant differences. Scale bar = 200 μm.

LPS applied at ZT6 did not significantly induce pGSK3β increase in the SCN at any time point, although on average, the level of pGSK3β in all LPS-treated sections was 1.4648 (± 0.1691) higher than the level in control sections (Fig. 5D, E). LPS applied at ZT15 significantly reduced the number of pGSK3β immunopositive cells in the vlSCN within 8 h (P = 0.0073) (Fig. 5D). There was no significant change in pGSK3β levels in dmSCN at night (Fig. 5E).

## Discussion

Several reports have focused on the effect of systemic inflammation on various aspects of the circadian physiology (Duhart et al., 2013, 2016; Guerrero-Vargas et al., 2014; Leone et al., 2006, 2012; Okada et al., 2008; Marpegán et al., 2005; Paladino et al., 2010, 2014). Most of these studies, however, only used one daytime period for inflammatory stimulation, although it is well understood that the susceptibility of organisms to inflammatory stimuli is affected by the circadian clock. Previously, we have shown that LPS applied either during the day or at night affects the phosphorylation of STAT3 in the SCN differently (Moravcová et al., 2016). In this study, we followed the changes in the *Stat3* mRNA and several selected markers that are known for their sensitivity to extra-clock events and may serve as probes to the main oscillatory loops and posttranslational events. The daytime point of infection coincides with high levels of STAT3 protein and of clock genes *Per1, Per2* and *Nr1d1* mRNAs (Moravcová et al., 2016; Shearman et al., 1997; Albrecht et al., 1997, Albrecht, 2012), and was chosen to deduce a possible LPS-induced decrease of their levels. The night point coincides with low levels of selected gene transcripts, which allows for measuring a possible induction in response to LPS. We used 1 mg/kg of LPS, which has been shown to suppress the expression of *Per1* gene in the SCN the day following the injection at ZT1 (Okada et al., 2008) and to induce the phosphorylation of STAT3 in the SCN astrocytes (Moravcová et al., 2016).

The behavioural effect of the 1 mg/kg LPS dose was controlled using locomotor activity monitoring. The maximal reduction of locomotion was observed the first day after the LPS treatment, as expected, but the gradual recovery process proceeded faster when the animals were infected during the day compared with at night. Total activity counts and the amplitude of the behavioural rhythm seemed to return to their original state faster after the daytime LPS injection. Considering the locomotor activity measurement as one of the markers of sickness behaviour, which typically occurs after infection (Hart, 1988), our observation suggests that the sickness behaviour after the LPS treatment can be improved more quickly if animals are infected during the day. This corresponded well to the organism’s reduced reaction to a parasitic infection when the organism was infected during the day compared to at night (Kiessling et al., 2017). It has been demonstrated that LPS administration at ZT14 triggers higher plasma level of cytokines IL-6 and TNF-α as compared with administration at ZT2, which has been associated with daily changes in SCN activity (Guerrero-Vargas et al., 2014). It may, therefore, be possible that higher levels of inflammatory cytokines at night decelerate the amelioration of sickness or worsen the symptoms.

STAT3 plays various roles in many cell types, but it is best known as a regulator of the inflammatory response and of cancer growth (Rummel, 2016; Yu et al., 2014). In the brain, STAT3 plays a significant role in astrocyte reactivity in response to pathological conditions in the nervous tissue (Ceyzériat et al., 2016). Our data showed that LPS administration at ZT15 induced about 10 times higher *Stat3* mRNA levels than at ZT6. There was also a significant difference in mRNA levels between day and night among the control animals, so we performed *in situ* hybridisation to detect the circadian profile of *Stat3* expression in the SCN. The data showed that spontaneous *Stat3* expression in the SCN was high during the day and low during the night. It is possible that LPS-induced upregulation may be limited by “ceiling” effect of the *Stat3* transcription which cannot be exceeded after the daytime injection. Yet, in average 15 times upregulation of *Stat3* mRNA above the baseline level at the time, when the STAT3 is not usually incorporated into the SCN signalling cascades, could serve as the specific time signal that may affect the production of cytokines and inflammatory response within the circadian clock.

It has been shown that STAT3 in the SCN is expressed exclusively in astrocytes (Moravcová et al., 2016). We can, therefore, assume that the rhythmic expression of its mRNA and transcriptional upregulation by the LPS also occur in astrocytes. Growing evidence supports the role of astrocytes in mediating the immune signals to the SCN (Leone et al., 2006; Duhart et al., 2013, 2016; Moravcová et al., 2016). The chemokine Ccl2, which is secreted in response to immune activation by SCN astrocytes *in vitro* (Duhart et al., 2013), has recently been shown to play a role in the circadian response to immune activation (Duhart et al., 2016). Moreover, the treatment of SCN astrocytes isolated *in vitro* with TNF-α modulates the clock genes expression of SCN astrocytes (Marpegán et al., 2005; Duhart et al., 2013). STAT3 can be part of the signalling by which the SCN astrocytes communicate the pathological conditions to the circadian clock. Interestingly, STAT3 has also been implicated in the regulation of sickness behaviour, including adipsia and febrile response to the LPS (Damm et al., 2013), which may support the significance of STAT3 signalling in the immune response of the hypothalamus.

Regulation of *Per* genes expression is the principal mechanism by which photic or nonphotic stimuli adjust the circadian phase to the external time (Albrecht et al., 1997; Akiyama et al., 1999). Several studies have reported decreases in *Per1* and *Per2* levels in response to various nonphotic stimuli when applied during the day (Maywood et al., 1999; Horikawa et al., 2000; Fukuhara et al., 2001). The first study concerning the effect of systemic LPS on *Per* gene expression in the SCN did not report any changes in either *Per* gene after 50 µg/kg of LPS was administered at ZT22, i.e. at the late night (Takahashi et al., 2001), which is similar to our observations after LPS at ZT15, i.e., at the early night. On the other hand, a 24-h treatment with the cytokine TNF-α suppressed the transcription of *Per1* and *Per2* in mice fibroblasts and livers (Cavadini et al., 2007), but the short-term stimulation led to upregulation of *Per1* and *Per2* in fibroblast culture (Petrzilka et al., 2009). Accordingly, we revealed a mild increase of *Per2* after 8 h and a decrease of *Per1* 24 h after the LPS was injected at ZT6 but not at ZT15. In the SCN, the prolonged 24-h responsiveness to daytime LPS application, as observed for *Per1* expression, has been reported before; under similar experimental conditions, upregulation of the p65 subunit of NF-κB transcription complex and immediate early gene *c-Fos* in the SCN was observed only 24 h after the LPS treatment (Beynon and Coogan, 2010). The suppression of clock gene expression in the SCN also occurred within the next circadian cycle after LPS administration (Okada et al., 2008). Furthermore, in our previous study, we observed a high level of phosphorylated STAT3 on Tyr705 after 24 h of LPS treatment during the daytime (Moravcová et al., 2016). Our findings on how LPS affected *Per* genes expression thus do not contradict previous observations and stress the significance of the 8-h and 24-h delays of the circadian clock’s responsiveness to inflammatory daytime stimulation.

The effect of LPS on *Nr1d1* expression in the SCN has not yet been studied. The significance of the protein product of this gene—nuclear receptor REV-ERBα—was demonstrated in macrophages, where REV-ERBα negatively regulates the inflammatory function by repressing IL-6 and chemokine Ccl2 gene induction following an LPS challenge (Sato et al., 2014a, b). Whether the upregulation of *Nr1d1* expression 8 h after the LPS injection affects the levels of IL-6 or Ccl2 in the SCN can be speculated upon. The Ccl2 in the SCN shows the circadian rhythm, with a high level during the night. The Ccl2 level already decreases by ZT23, the time of *Nr1d1* upregulation after early-night LPS injection (Duhart et al., 2016). Considering that Ccl2 can be induced within 1 h after the LPS treatment, the delayed upregulation of REV-ERBα may enforce the clock-driven suppression of the Ccl2. A similar mechanism has been proposed for the role of REV-ERBα in macrophages (Gibbs et al., 2012).

The pERK1/2 level in the SCN is rhythmic and oscillates in antiphase between vlSCN and dmSCN (Pačesová et al., 2015). Its level in the vlSCN can be induced within minutes by the light pulses at night (Obrietan et al., 1998). Our data showed pERK1/2 upregulation within 2 h after LPS treatment at ZT6 in both parts of the SCN, and downregulation of its high level in the vlSCN 8 h after the LPS injection at ZT15. Besides the well-documented LPS-induced pERK1/2 level in macrophages and *in vitro* osteoblasts (Rawadi et al., 1998; Chen and Wang, 1999; Daigang et al., 2016), the dramatic increase in pERK1/2 immunoreactivity was also apparent 2 h after LPS was administered in the paraventricular nucleus of the hypothalamus (Singru et al., 2008). Two studies observed the reduction of a spontaneously high pERK1/2 level in the vlSCN at night: it declined within 2 h after light pulses at night (Červená et al., 2015) and after an opioid challenge at night (Pačesová et al., 2015).

Similarly to pERK1/2, the high level of pGSK3β in the vlSCN was reduced 8 h after LPS was administered at night. GSK3β promotes the nuclear translocation of clock protein PER2 (Iitaka et al., 2005) and triggers the proteasomal degradation of CRY2 (Kurabayashi et al., 2006). It has been demonstrated that GSK3β promotes the LPS-induced production of proinflammatory cytokines in the microglial cells, possibly in cooperation with STAT3 (Beurel and Jope, 2009; Green and Nolan, 2012). It may be important to reiterate that GSK3β kinase is active in its dephosphorylated state. It is, therefore, possible that a short-term increase in its activity contributes to LPS-induced changes in cytokine level at night. In the circadian pacemaker, the transient GSK3β-induced increase of phosphorylation of clock proteins could change the dynamics of the clockwork mechanism and thus participate in maintaining a steady-state clock status.

The induction and reduction of the clock genes and pERK1/2 with the pGSK3β levels in the SCN in our study seemed to be relatively mild. Although significant, the induction does not vary by order, such as after the light stimulus, for example. A majority of the studies concerning the effect of LPS or cytokines on the SCN markers have shown similar magnitudes (Beynon and Coogan, 2010; Paladino et al., 2014; Moravcová et al., 2016). Because different doses of LPS and cytokines were used in these studies, the magnitude of the observed changes does not seem to depend on the intensity or specificity of the inflammatory stimuli. These alterations could reflect the hypothalamic homeostatic drive, which helps the circadian clock to cope with the acute pathological environment. We observed the quick response of *Stat3* and pERK1/2, which reacted within two hours, and—together with other authors—the delayed response of *Per1* gene after the daytime stimulus. The most active period of LPS-induced changes in SCN state seemed to occur about 8 hours after the infection; at this time, the *Per2* was induced after the daytime LPS injection, compared to *Nr1d1* after the night-time LPS injection. Regarding the clockwork machinery, both effects should result in phase shifts of the circadian oscillations. In addition, the increase of active GSK3β may lead to advances of the clock phase (Osland et al., 2011). It is possible that the mild processes within the circadian clockwork help to balance the possible dysregulation of the circadian clock output.

## Conclusions

The present study shows that the time of LPS administration affects the recovery rate of locomotor activity rhythm and induces the transient changes in clock gene expression and the levels of pERK1/2 and pGSK3β in the rat SCN that may be a part of the steady-state function of the clock in mild pathological conditions. We also provide the first report on the circadian rhythmicity of *Stat3* gene expression in the SCN, and we demonstrate that the LPS administration induces not only phosphorylation of STAT3 that has been shown previously, but also its transcription and regulate thus significantly *Stat3* mRNA level in the SCN.

## Acknowledgments

We thank Dr. Peter Ergang for his help with the laser dissection. This work was supported by Charles University Grant Agency no. 361115; by the Czech Science Foundation, contract grant number 18- 08423S, and by project no. LO1611 with financial support from the MEYS under the NPU I programme.

